# Body mass index, earnings and partnership: genetic instrumental variable analysis in two nationally representative UK samples

**DOI:** 10.1101/608588

**Authors:** Amanda Hughes, Yanchun Bao, Melissa Smart, Meena Kumari

## Abstract

In high-income countries there is an established link between high body mass index (BMI) and low income, but the direction of this association is unclear. Recent analyses in a large UK population using genetically-instrumented BMI supported a causal influence of BMI on household income, educational attainment and job class. Since analyses were based on an age-restricted and relatively wealthy population, it is unclear whether results are generalizable, and limited income data precluded decomposition of household income effects into own-income and partnership effects. Investigation is therefore warranted in more representative UK populations where associations may differ, and where individual and partner-based mechanisms can be studied separately.

Data came from two nationally-representative samples, the UK Household Longitudinal Survey (UKHLS) and the English Longitudinal Study of Ageing (ELSA). Analysis was conducted in each sample, with results then pooled by meta-analysis. We used externally-weighted polygenic scores based on the latest genome-wide association study for BMI to examine the influence of genetically-instrumented BMI on earnings, probability of employment, job class conditional on working, likelihood of partnership, and partner’s earnings.

A one-unit (kg/m^2^) increase in genetically-instrumented BMI was associated with a roughly 9% decrease in own monthly earnings (pooled coefficient: 0.91, CI:0.86, 0.97) and lower probability of employment (OR: 0.89, CI:0.83, 0.96) or having a university degree (OR: 0.95, CI:0.90, 0.99). Employed individuals with higher genetically-instrumented BMI were less likely to have professional or managerial occupations (OR: 0.91, CI:0.86, 0.96). No associations were seen with partnership. A one-unit increase in BMI was associated with a 5% decrease in partners’ earnings, but estimates were imprecise (pooled coefficient: 0.95, CI:0.88,1.01).

Results are consistent with a negative influence of body mass index on a range of labour market and educational outcomes for both men and women.

**Key Messages:**

- Higher genetically-instrumented BMI was associated with lower earnings and odds of working
- Also with lower odds of holding a managerial/professional occupation or a degree
- No associations were seen with probability of cohabiting partnership

## Background

An association of disadvantaged socioeconomic position (SEP) and high BMI is widely documented in high-income countries(1, 2), but the mechanisms involved are poorly understood. Firstly, low income may directly impact on body weight through diet, with calorie-dense, nutrient-poor foods often chosen to stretch a restricted food budget(3). Secondly, the relationship of adult SEP and adiposity may be influenced by common factors, for example by childhood socioeconomic position (4, 5). Lastly, adiposity may influence SEP. Employer discrimination against heavier candidates may reduce opportunities for employment or promotion(6), and impact of obesity-associated health problems may affect an individual’s labour market success(7). A pathway via educational attainment, which may also be influenced by BMI(8), is also possible. Disentangling causality in the relationship of adiposity and SEP is therefore challenging, but Mendelian Randomization (MR) - use of genetic variants as instrumental variables - provides an opportunity to investigate the causal influence of BMI on SEP. While BMI and SEP can both change across a person’s life in response to environmental influences, gene variants do not, and gene variants associated with BMI may be used as instruments for BMI robust to reverse causation and classical sources of confounding(9). Two recent papers have taken this approach using data from the UK Biobank(10, 11). The first used data from the pilot sample and supported a causal association of BMI on household income only for women. The second, using data from the later release of over 350,000 individuals, reported a causal negative association of higher BMI for both women and men on household income, educational attainment and probability of having a skilled occupation(11). Impact of BMI on partnership status differed by gender: high BMI lowered probability of partnership for women, but low BMI lowered probability of partnership for men. However with an approximately 5% response rate in UK Biobank, generalizability of findings from this highly selected population has been called into question(12, 13), warranting further investigation in more socioeconomically representative UK samples. Investigation of income was also limited by the measures in UK Biobank: participants reported only total income household income, chosen from wide categories (e.g. <£18,000, £18,000-£30,999). Thus, estimates are affected by income of other individuals in the household, non-labour income sources, and loss of information by categorization. While mechanisms such as weight-related stigma are purported to act on labour income specifically, estimates in UK Biobank include other income sources which may be unaffected or increased by a high BMI. A separate recent study in 2,064 working-age Finnish individuals reported a negative influence of genetically-instrumented BMI on earnings and employment status, but a positive influence on social benefits income(14). However, despite the more representative sample and high-quality income data from administrative records, the small sample size precluded gender-stratified analysis, and results were sensitive to specification.

In the UK Household Longitudinal Study (UKHLS) and the English Longitudinal Study of Ageing (ELSA) – two studies with extensive information on labour and non-labour income of participants and their partners - we examine the relationship of genetically-instrumented BMI with earnings of individuals and their partners. Since observational BMI-SEP associations have been shown to differ by gender(1, 15), we conduct separate analyses for men and women. We specifically consider earnings from employment or self-employment, since the principal mechanisms proposed for a causal effect (discrimination by employers and health-related impact on productivity) are purported to impact on labour income specifically. We examine separately relationships with own and partners’ earnings, to investigate whether previously reported associations of BMI and household income in UK Biobank are explained by own labour market success or partnering mechanisms. A link between genetically-instrumented BMI and partners’ earnings could occur for several reasons: cross-trait selection pairing for instance thinner women with higher-earnings partners, single-trait selection pairing heavier individuals together, with both partners experiencing negative influence of own BMI on earnings, or an influence of one partners’ genotype on the other’s phenotype via environmental pathways. Although smaller than UK Biobank, the combined sample of 9447 was several times larger than used in the Finnish study(14). Unlike either study, we use genetic instruments based on the largest and most recent GWAS of BMI, which explain substantially more variation in BMI than SNPs identified in earlier GWAS(16).

## Methods

### Study participants

The UKHLS is an annual longitudinal survey of over UK 40,000 households. It consists of a larger General Population Sample (GPS), a stratified clustered random sample of households representative of the UK population which joined in 2009-10, and a smaller component from the pre-existing British Household Panel Survey (BHPS)(17). Blood samples taken during a nurse visit approximately five months after the main wave 2 interview (GPS participants) or wave 3 interview (BHPS participants), with eligibility criteria detailed elsewhere(18). 9944 individuals were genotyped, restricted to those of white ethnicity to avoid population stratification. Individuals aged <25 were excluded, since below 25 low earnings may reflect individuals in higher education; individuals aged >73 were excluded, given the very low rate of labour market participation beyond this point, and for consistency with previous UK-based analyses (10, 11). Analyses of partner’s earnings were restricted to individuals whose partners were also aged 25-73. For pairs of individuals related by r2=0.20 or more one was randomly chosen for exclusion. Exclusion for missing covariates and zero–value inverse-probability weights left a final sample size of 6,785 for analyses of individual earnings and 4,898 for analyses of partner’s income.

The English Longitudinal Study of Ageing began in 2001 as a sample of English adults aged 50+. So that earnings data was measured in a comparable economic climate for ELSA and UKHLS participants, ELSA data was drawn from the wave 6 main and biomedical visits, which took place in 2012-13. Genetic information was available for 5,260 individuals of white ethnicity who took part at wave 6 and were not related, again filtering at r2=0.20. Analysis was similarly restricted to participants aged 73 or younger, and for analyses of partner’s earnings, to individuals with partners aged 73. Exclusion for missing data on BMI, earnings and covariates left a final sample size of 2662 for analyses of individual earnings and 2000 for analyses of partner’s income.

## Measures

### Earnings of individuals and their partners

In both surveys, outcomes were defined based on participant-reported data on income from employment and self-employment. We consider individual earnings rather than total or household income, as the mechanisms under study relate to how much a person can earn, or how much others are willing to pay them, rather than the economic resources at their disposal. For UKHLS participants we consider monthly earnings at the annual interview corresponding to the nurse visit (W2 for GPS participants, W3 for BHPS participants). This incorporates income from main employment, self-employment, and second jobs. In ELSA, we consider monthly earnings at the wave 6 interview, incorporating net income from main employment, self-employment, and second jobs. We use net earnings, since in ELSA comparable-quality data for gross earnings was not available: analyses for UKHLS using gross earnings are presented in supplementary tables. To reduce influence of outliers and remove likely errors, the top 0.5% of observations were removed, with remaining observations log-transformed. Partnership status was defined by cohabitation. Data on partners’ earnings was available for 98.1% of participants cohabiting with a partner in UKHLS, and 97.9% of these participants in ELSA.

### Employment status, occupational class

Self-reported employment status was dichotomized as in work (employment or self-employment) or not in work. This group contained participants who were retired, jobseekers, homemakers, out of the labour force due to ill-health, or in any other category besides employment or self-employment. Within working individuals, influence was explored of BMI on likelihood of holding a professional or managerial profession, using a binary measure comprising NS-SEC groups 1 and 2 as opposed to all other groups.

### Polygenic score

ELSA participants were genotyped using Illumina Omni 2.5M array, and UKHLS participants using the HumancoreExome array. In both surveys, imputation was carried out for SNPs with minor allele frequency of >1% using Minimac to the European component of 1000 Genomes. For both surveys, a polygenic score (PGS) was calculated based on the most recent and largest genome-wide association study of anthropometric traits(16). An externally-weighted PGS for each participant was derived by summing their number of BMI-increasing alleles, with each allele weighted by the effect size of the SNP-BMI association in the GWAS. Use of a summary score reduces bias compared to including SNPs as individual instruments, while external weighting increases power because different SNPs can have considerably different effects on the instrumented variable(19). From full GWAS data, SNPs associated with BMI at p<5×10^−8^ which were available in UKHLS and ELSA and passed quality controls were identified, and resulting SNPs were clumped in MR-Base(20) at r^2^=0.001. This resulted in a 480-SNP PGS in UKHLS and a 518-SNP PGS in ELSA (Supplementary Tables 2-3).

### Height, weight and confounders

In both surveys, BMI was calculated from height and weight measured by a nurse, using a portable stadiometer and digital floor scales. Participants gave estimated weights if heavier than 130kg, where the scales become inaccurate(18). In all analyses the first 10 genetic ancestry principal components were included to account for possible population stratification. Partnership was defined by self-reported cohabitation with a partner (no/yes). Age, gender and educational qualifications came from questionnaire information. This was first categorised into no qualifications/qualifications below degree/university degree or equivalent, and then standardized by gender and 5-year age band to take account of generational differences in the distribution of education, following a procedure detailed elsewhere (21). This resulted in a score between 0 and 1, with higher scores indicating more education relative to peers. Participants rated their overall health as excellent/very good/good/fair/poor, again analysed as continuous. Employment status, based on participants own description of their current situation, was dichotomised to employed or self-employed/other. UK Government office region (GOR) was identified from participant postcodes and classified as North East/North West/Yorkshire and Humberside/East Midlands/West Midlands/East Anglia/London/South East/South West/Scotland/Wales. Smoking status was categorised as never smoker, ex-smoker, and current smoker.

## Analysis

All models were inverse-probability weighted using blood weights from the cross-wave nurse visit (UKHLS) and the wave 6 nurse visit (ELSA) to address non-response and sampling bias. Since both samples are complex survey data, STATA’s *svyset* command was used in all regressions to account for sample stratification and clustering, declaring household IDs as the primary sampling unit. Observational models used linear and logistic regression for continuous and binary outcomes respectively. Instrumented models used linear two-stage least squares regression for instrumental variables (STATA’s *ivregress*, which is compatible with survey data). Following previous analyses of this question(10, 11), IV estimates for binary outcomes were based on manual two-stage regression, with predicted BMI based on the PGS entered into a second-stage logistic regression model. IV models adjust for gender, age and age^2^, and the first ten principal components only: with a valid instrument, adjustment for confounders of observational associations is not necessary, and can introduce bias if those factors were not included in the GWAS used as the source of the gene-exposure associations(22). We therefore include only in observational models factors likely to confound observational associations: Government Office Region within the UK, by which both average BMI(23) and average earnings vary(24), smoking, self-rated health, and educational qualifications. To facilitate comparison, figures additionally show OLS associations adjusted for the same factors as IV models (age, age^2^, gender and principal ancestry components only). Analyses were conducted for all participants, and for men and women separately. Coefficients from the two surveys were pooled using STATA’s metan package, with both fixed and random pooled effects presented.

### Robustness checks

Within each survey, standard robustness checks for instrumental variable analysis were performed. In UKHLS, where comparable-quality data on gross earnings was available, models were repeated using gross earnings. To assess possible directional pleiotropy (failure of the exclusion restriction due to multiple functions of SNPs), we compared results from inverse-variance weighted (IVW), MR-Egger, and MR-Median regression (25, 26).

## Results

### Sample description

UKHLS participants were younger than ELSA participants, with a mean age of 51.9(12.8), vs 64.9(4.6). Reflecting age differences in the populations, many more UKHLS participants were in work (62.5% vs 30.3%), and labour income was substantially higher for UKHLS participants, and their partners (Table 1). UKHLS participants were considerably more educated (e.g., 36.3% with a degree, vs 23.0%), but self-rated health was similar between the studies, with the proportion of participants reporting only fair or poor health 19.5% in UKHLS, and 19.0% in ELSA.

**Table 1:** Descriptive Characteristics of Analytic Sample.

| Table 1: Descriptive Characteristics of Analytic Sample |  |  |  |  |  |  |  |
| --- | --- | --- | --- | --- | --- | --- | --- |
|  |  | UK Household Longitudinal Study, 2010-12 |  |  | English Longitudinal Study of Ageing, 2012-13 |  |  |
|  |  | All<br>N=6785 | Men<br>2,931 | Women<br>3,854 | All<br>2662 | Men<br>1220 | Women<br>1442 |
|  |  | *Mean (SD)* | *Mean (SD)* | *Mean (SD)* | *Mean (SD)* | *Mean (SD)* | *Mean (SD)* |
| Age(y) |  | 51.9(12.8) | 52.6(12.7) | 51.3(12.8) | 64.9(4.6) | 65.0(4.6) | 64.9(4.6) |
| BMI (Kg/m<sup>2</sup>) |  | 28.3(5.4) | 28.5(4.7) | 28.2(6.0) | 28.2(5.1) | 28.3(4.6) | 28.2(5.6) |
| Monthly earnings (£) |  | 899.8(984.9) | 1183.4(1119.5) | 684.1(804.5) | 410(782) | 575(924) | 271(605) |
| Partner's monthly earnings (£) |  | 1010.9(1063.0)<sup>b</sup> | 753.2(850.6)<sup>c</sup> | 1236.6(1173.6)<sup>d</sup> | 470(842)<sup>e</sup> | 383(677)<sup>f</sup> | 556(970)<sup>g</sup> |
|  |  | % | % | % | % | % | % |
| Employment status | Employed/self-employed | 62.5 | 67.7 | 58.6 | 30.3 | 35.9 | 25.5 |
|  | Not in work | 37.5 | 32.3 | 41.4 | 69.7 | 64.1 | 74.5 |
| Occupational social class (NS-SEC 2010) | Professional/managerial | 26.5 | 29.3 | 24.3 | 11.4 | 14.2 | 9.0 |
|  | Other | 35.3 | 37.6 | 33.6 | 18.9 | 21.7 | 16.5 |
|  | Not in work | 37.5 | 32.3 | 41.4 | 69.7 | 64.1 | 74.5 |
|  | Missing | 0.7 | 0.8 | 0.7 |  |  |  |
| Highest Educational Qualification | Degree | 36.3 | 36.3 | 36.3 | 23.0 | 28.4 | 18.4 |
|  | A-level or equivalent | 19.1 | 22.8 | 16.3 | 29.0 | 32.5 | 25.9 |
|  | GCSE or equivalent | 21.4 | 19.3 | 23.0 | 23.9 | 21.0 | 26.3 |
|  | Foreign/other | 11.0 | 11.3 | 10.7 | 6.2 | 3.6 | 8.5 |
|  | No qualifications | 11.9 | 10.0 | 13.3 | 17.6 | 14.1 | 20.5 |
|  | Missing | 0.4 | 0.3 | 0.4 | 0.4 | 0.4 | 0.4 |
| Self-rated health | Excellent/very good/good | 80.4 | 80.3 | 80.5 | 80.9 | 79.4 | 80.2 |
|  | Fair/poor | 19.5 | 19.7 | 19.5 | 19.0 | 20.6 | 19.8 |
|  | Missing | 0.0 |  | 0.1 |  |  |  |
| Partnership status | No cohabiting partner | 28.3 | 22.9 | 32.4 | 76.6 | 82.3 | 71.7 |
|  | Cohabiting partner | 71.7 | 77.1 | 67.6 | 23.4 | 17.7 | 28.3 |
| <sup>a</sup>Externally weighted using beta coefficients from Yengo et al 2018. <sup>b</sup>N=4841 <sup>c</sup>N=2261 <sup>d</sup>N=2,580 <sup>e</sup>N=1,989 <sup>f</sup>N=987 <sup>g</sup>N=1002 |  |  |  |  |  |  |  |

### BMI and own earnings, employment status and occupational class

Figure 1 shows associations of per-unit (kg/m^2^) change in BMI with own log-transformed monthly earnings. In minimally-adjusted OLS models, a unit increase in BMI was associated with a 4% decrease in own earnings for all participants (exponentiated coefficients from pooled models: 0.96, CI:0.95, 0.98) and for women (pooled coefficient: 0.95, CI:0.94,0.97) but not men. Associations disappeared with adjustment for smoking, Government Office Region, and educational qualifications. Pooled IV estimates adjusted for age, age squared and principal components were less precise, but showed significantly negative association for all participants, corresponding to a roughly 9% decrease in earnings per unit change in BMI (exponentiated coefficient: 0.91, CI:0.86,0.97) and men (0.89, CI:0.80,0.98) with a weaker association in women (0.93, CI:0.86,1.01). Results for probability of working showed a very similar pattern. Significant negative associations for all participants and women in minimally-adjusted observational models (OR:0.97, CI:0.96,0.99, OR:0.96, CI:0.95,0.98) attenuated with adjustment for observational confounders. IV models showed significant negative associations for all participants (OR:0.94, CI:0.89,0.99), with similar associations in gender-stratified analyses. Among employed participants, IV models and minimally-adjusted observational models did show associations of BMI and reduced odds of holding a managerial or professional position which were similar by gender (IV estimate for all participants: OR:0.91, CI:0.86, 0.96).

**Figure 1:**
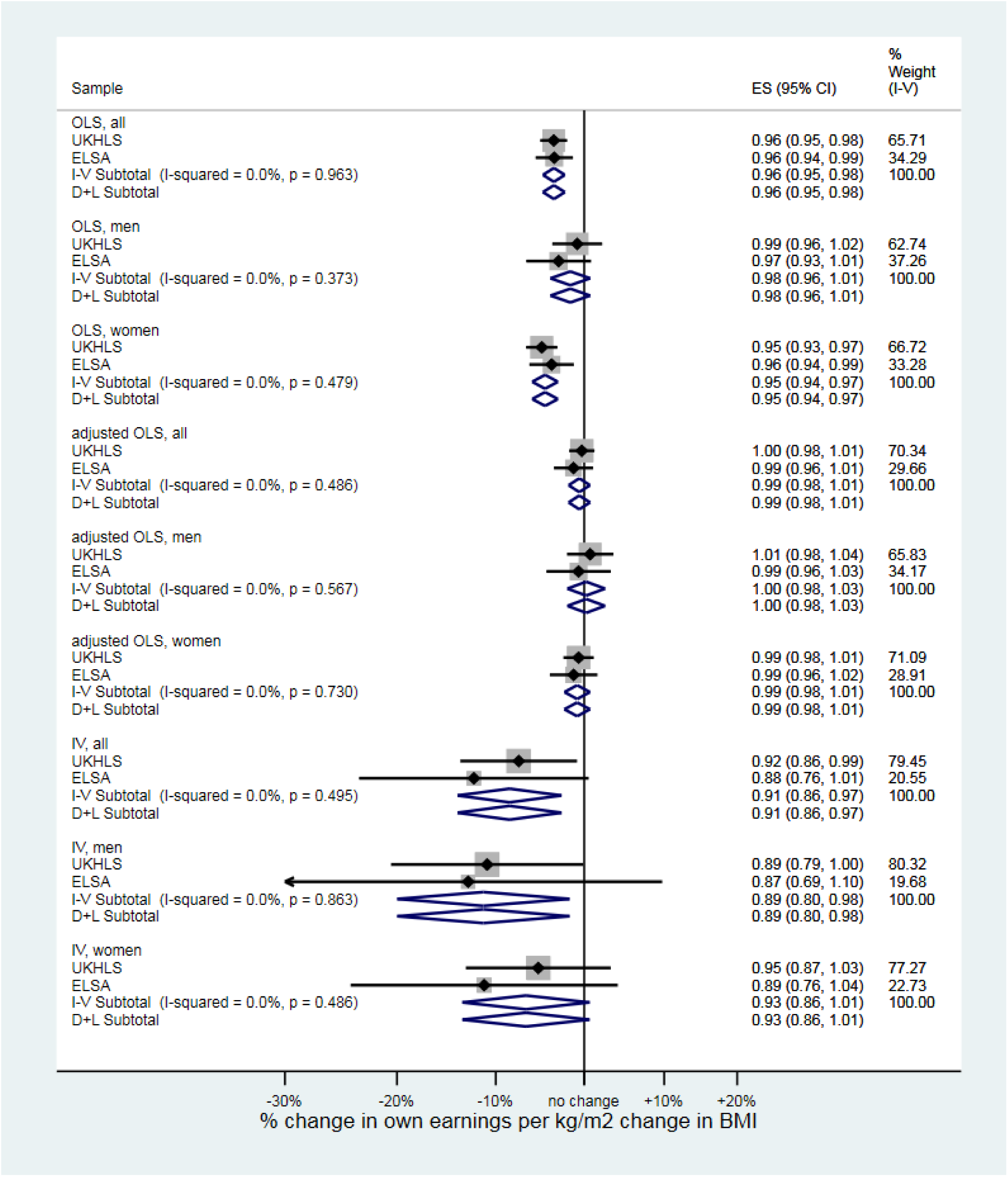
Associations of BMI and own earnings in UKHLS and ELSA. All participants, men and women.

**Figure 2:**
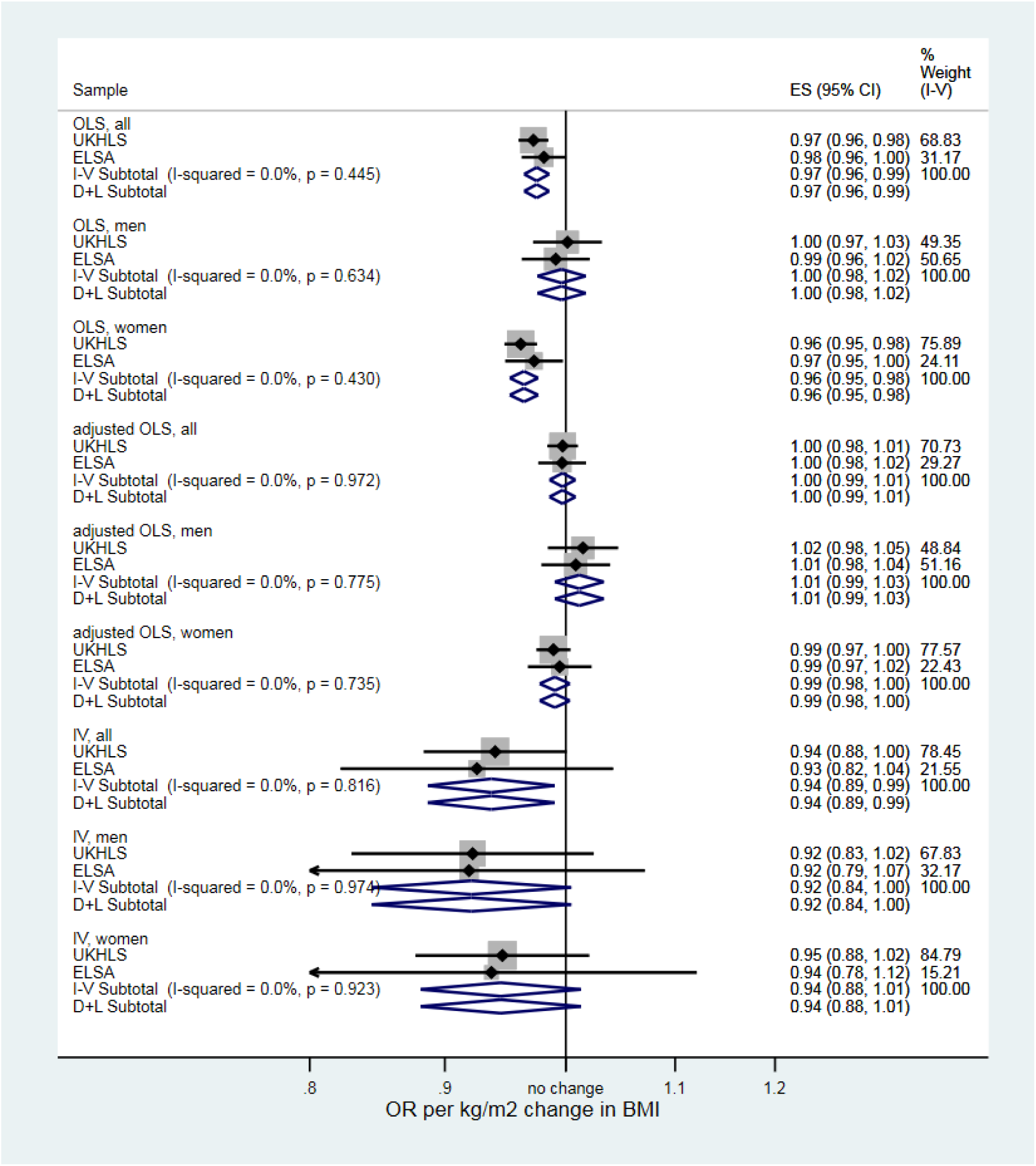
Associations of BMI and probability of working in UKHLS and ELSA. All participants, men and women.

**Figure 3:**
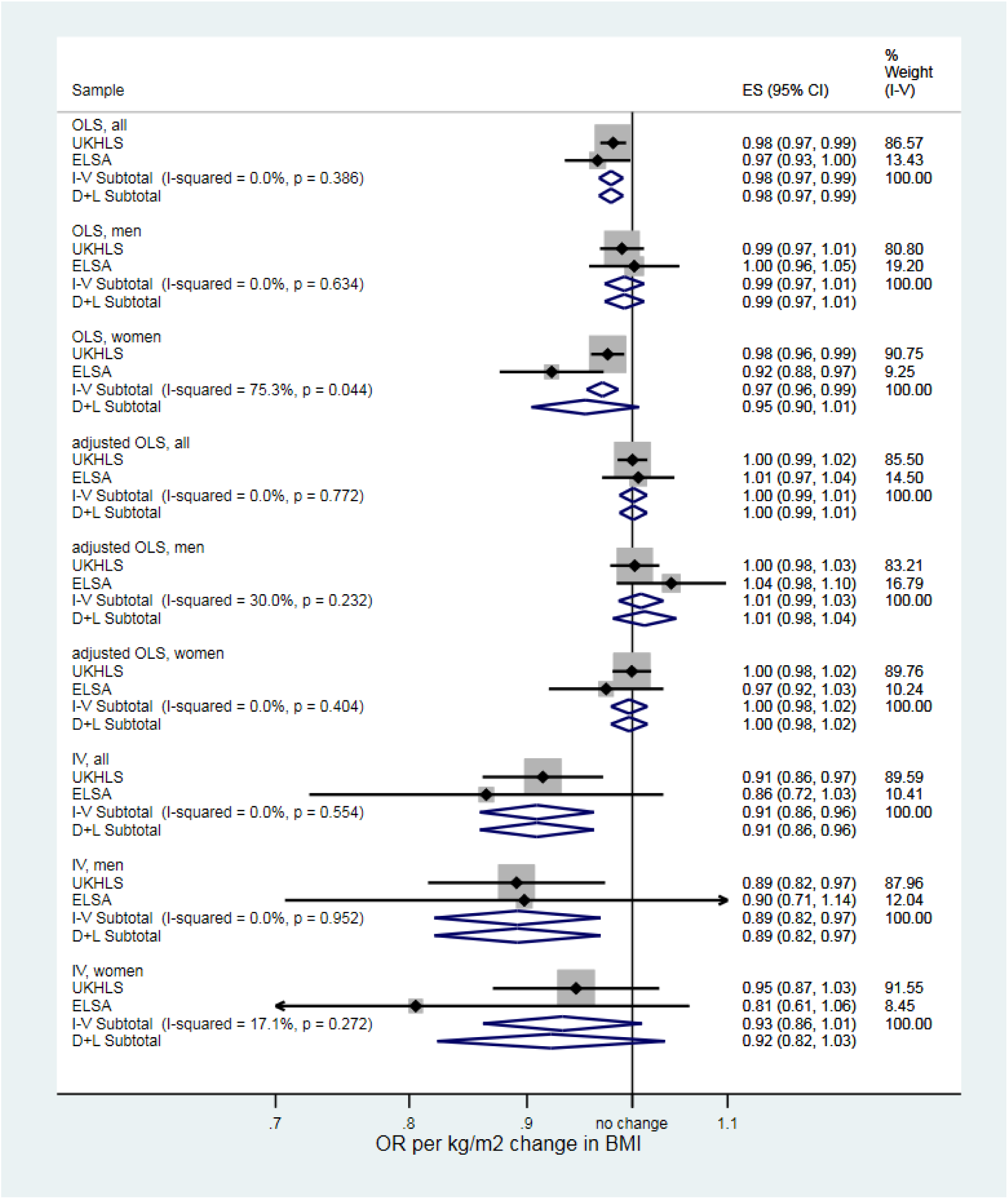
Associations of BMI and probability of managerial or professional occupation, conditional on current employment, in UKHLS and ELSA. All participants, men and women

### BMI, partnership and partners’ earnings

In OLS models, no associations were seen of women’s BMI and probability of cohabiting partnership (Figure 4), while men’s BMI was positively associated with partnership (pooled ORs for minimally-adjusted and fully-adjusted models: 1.02, CI:1.00,1.04 and 1.03, CI:1.01,1.05). In IV models, there was no evidence of associations for any group. In observational models of partners’ earnings for partnered individuals, negative associations were seen for all participants and for women but not men. In IV models, estimates were negative but imprecise, indicating across all participants a roughly 5% decrease in partner’s earnings per kg/m^2^ of own BMI (exponentiated coefficient: 0.95, CI:0.88,1.01). There was no evidence of gender modification.

**Figure 4:**
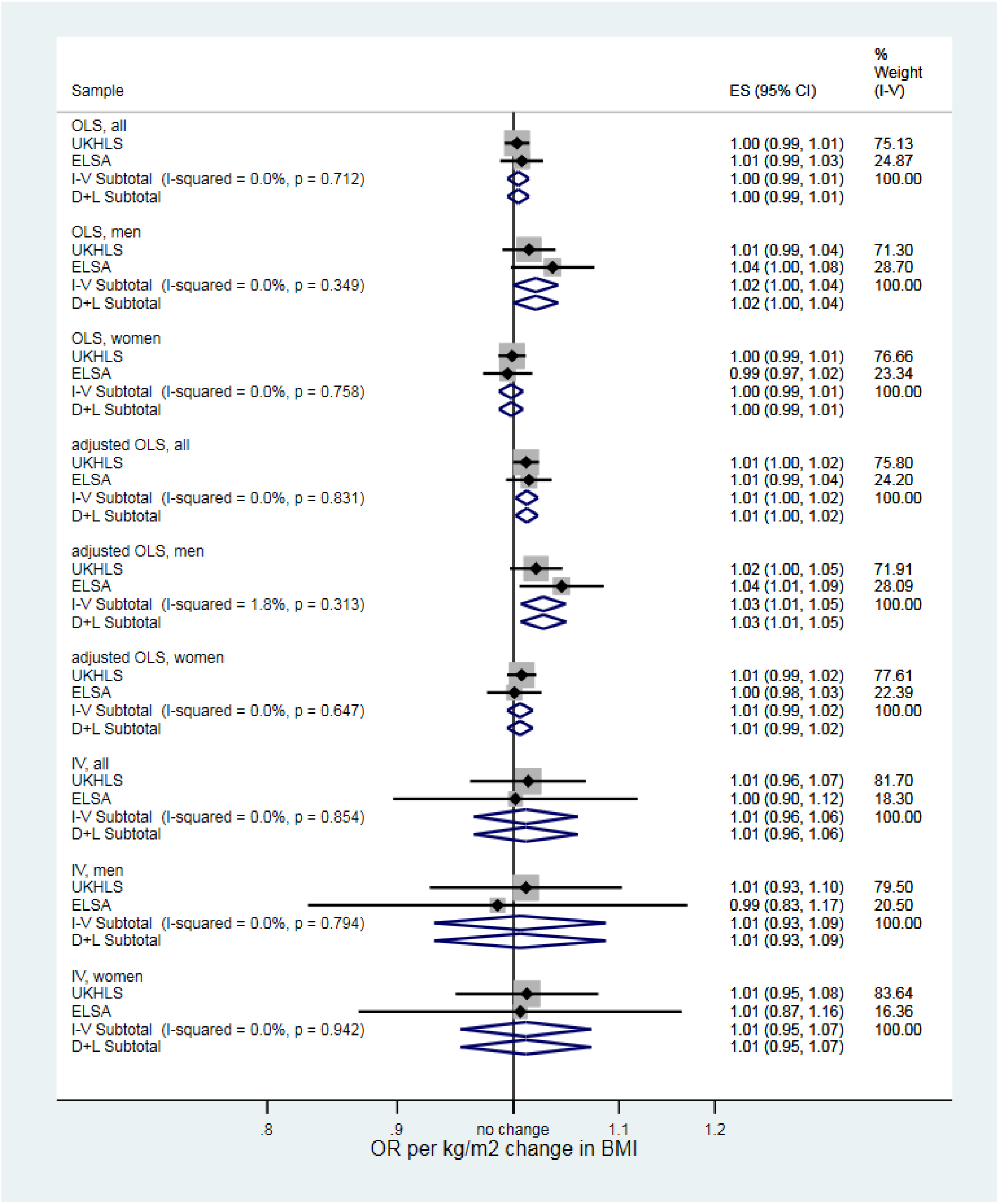
Associations of BMI and probability of cohabiting partnership in UKHLS and ELSA. All participants, men and women.

**Figure 5:**
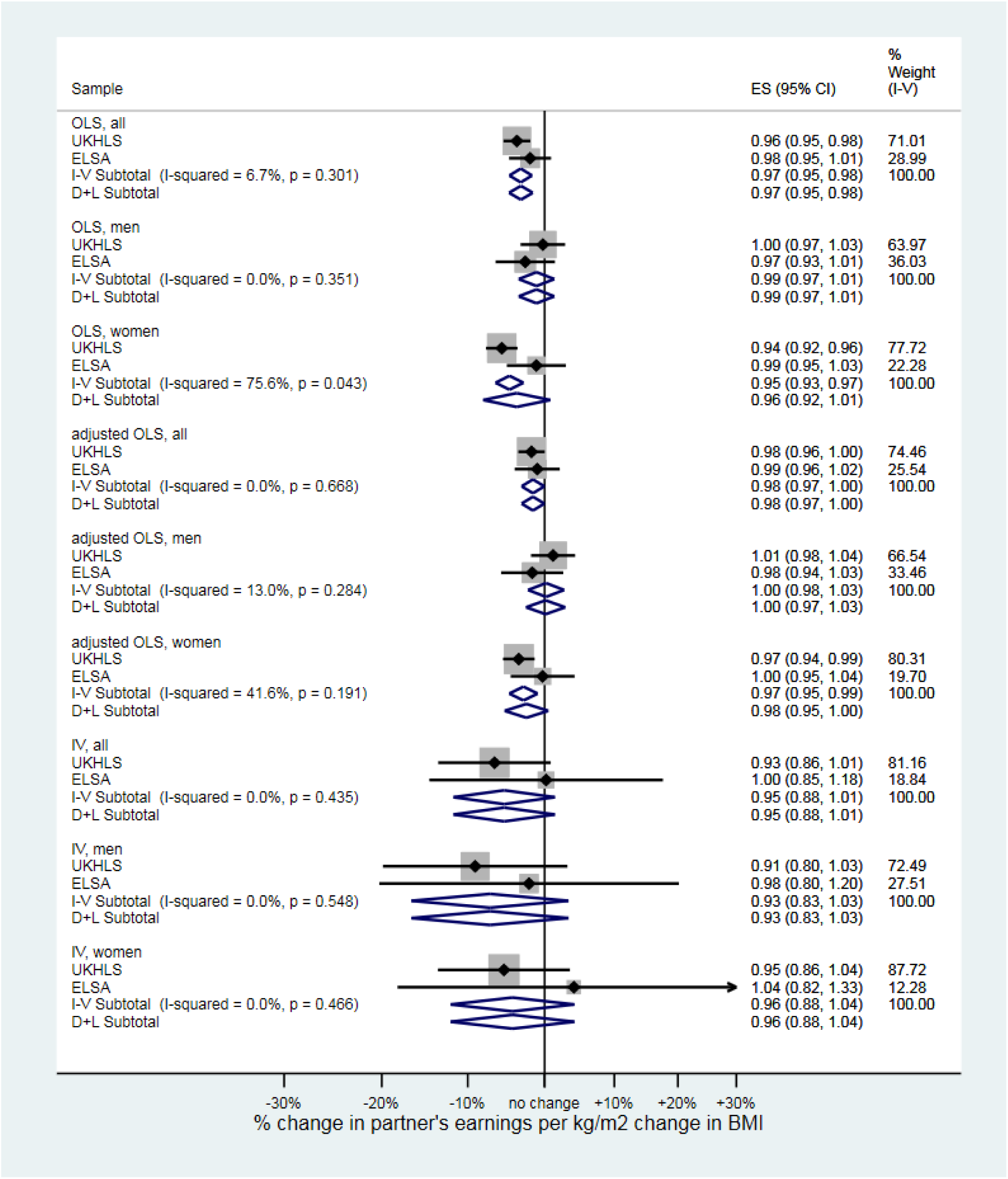
Associations of BMI and partner’s earnings in UKHLS and ELSA. All participants, men and women.

**Figure 6:**
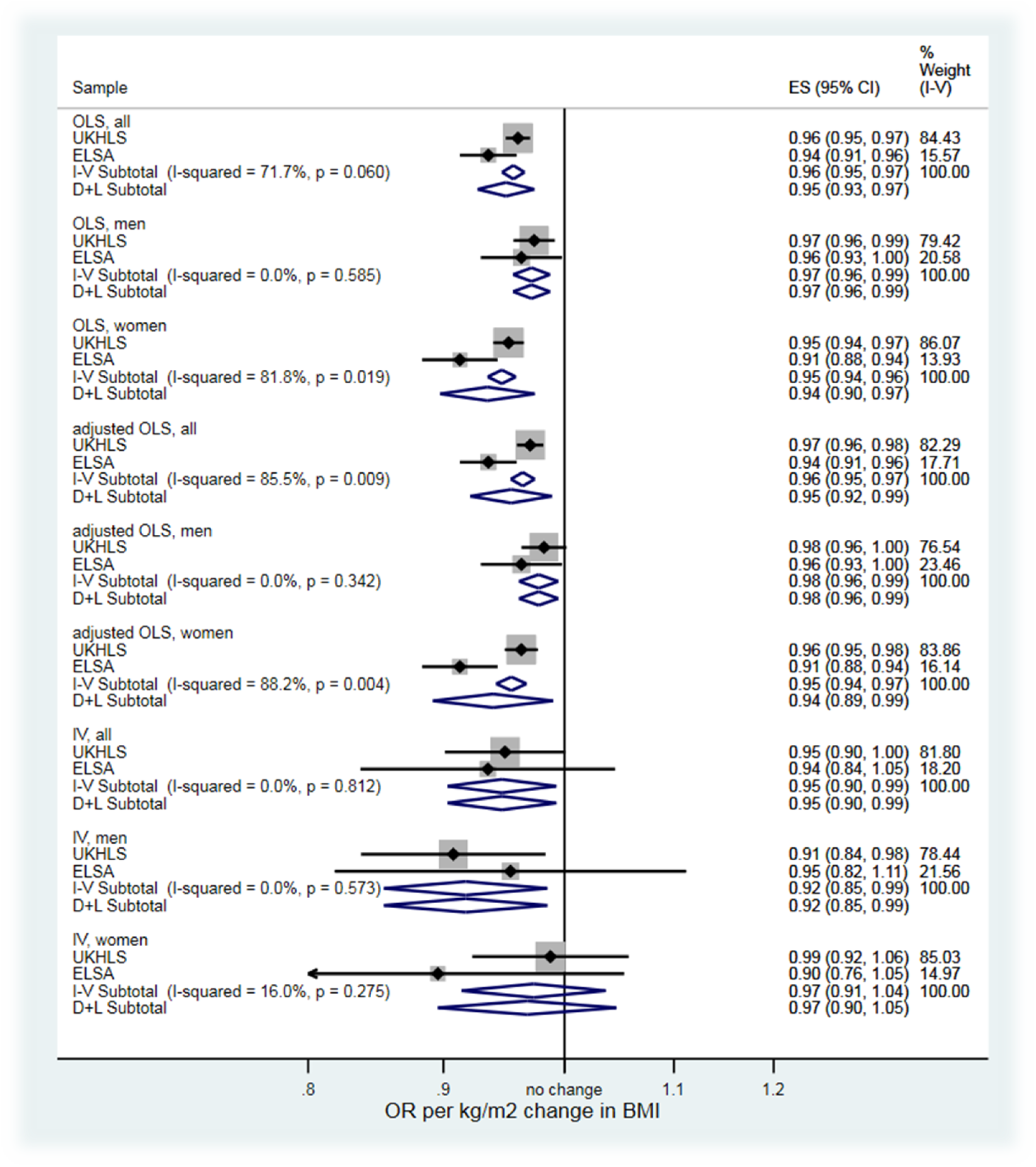
Associations of BMI and probability of having a university degree in UKHLS and ELSA. All participants, men and women

### BMI and Education: Probability of Degree

In OLS models, higher BMI was associated with lower odds of having a university degree for all participants, men and women. Associations were robust to adjustment for region and smoking status, but showed study heterogeneity, with stronger effects in ELSA. Random-effects pooled estimates with full adjustment were OR:0.96 (CI:0.92,0.99) for all participants, OR:0.98 (CI:0.96,0.99) for men and OR:0.94 (CI:0.89,0.99) for women. IV models showed negative effects without significant heterogeneity: pooled estimates were for all participants OR:0.95 (CI:0.90,0.99), for men OR:0.92 (CI:0.85,0.99) and for women OR:0.97 (CI:0.91,1.04) (fixed) and OR:0.97 (CI:0.90,1.05) (random).

### Association of the PGS with other relevant factors

In both surveys the PGS predicted worse self-rated health, especially for women (Table 2). There was evidence of associations with ever and current smoking for all participants and women in UKHLS, but not in ELSA (Table 2). In UKHLS, adjusted for age, age squared, gender and principal components, there was suggestive evidence that one partners’ standardized PGS predicted the other’s (beta: 0.05, CI:-0.00,0.11, p=0.06, N=6315). One partner’s instrumented BMI predicted the measured BMI of the other for all participants (0.21, CI:0.05,0.37, p=0.01, N=4,146) and women (0.30, CI:0.11,0.49, p=0.002, N=2,042) but not men (0.10, CI:-0.18,0.37, p=0.49, N=2,104). In ELSA, one partners’ PGS did not predict the other’s (beta: −0.01, CI:-0.08,0.07, p=0.88, N=1,312), and only women’s instrumented BMI predicted BMI of partners (beta: 0.41, CI:0.01,0.81, p=0.05, N=632).

**Table 2:** Associations of the PGS (per SD) with other relevant factors.

| **Table 2: Associations of the PGS (per SD) with other relevant factors** |  |  |  |  |  |  |  |  |
| --- | --- | --- | --- | --- | --- | --- | --- | --- |
| **UKHLS: (N=6785)** |  |  |  |  | **ELSA: (N=2662)** |  |  |  |
|  |  | Coef | CI | p |  | Coef | CI | p |
| Self-rated health | All | 0.10 | 0.07 - 0.12 | 0.00 | All | 0.07 | 0.02 - 0.11 | 0.005 |
|  | Men | 0.08 | 0.04 - 0.12 | 0.00 | Men | 0.03 | -0.03 - 0.10 | 0.34 |
|  | Women | 0.11 | 0.07 - 0.15 | 0.00 | Women | 0.10 | 0.04 - 0.16 | 0.002 |
| Height (cm) | All | -0.14 | -0.30 - 0.02 | 0.09 | All | -0.28 | -0.55 - -0.02 | 0.04 |
|  | Men | -0.25 | -0.51 - 0.00 | 0.05 | Men | -0.08 | -0.49 - 0.33 | 0.71 |
|  | Women | -0.05 | -0.26 - 0.16 | 0.65 | Women | -0.50 | -0.84 - -0.16 | 0.004 |
| Ever smoking | All | 0.05 | 0.02 - 0.08 | 0.00 | All | 0.01 | -0.04 - 0.06 | 0.74 |
|  | Men | 0.06 | 0.01 - 0.11 | 0.02 | Men | 0.04 | -0.04 - 0.12 | 0.32 |
|  | Women | 0.05 | 0.00 - 0.09 | 0.03 | Women | -0.02 | -0.09 - 0.05 | 0.62 |
| Current smoking | All | 0.07 | 0.03 - 0.11 | 0.001 | All | 0.03 | -0.04 - 0.10 | 0.43 |
|  | Men | 0.04 | -0.02 - 0.10 | 0.22 | Men | 0.07 | -0.04 - 0.18 | 0.21 |
|  | Women | 0.10 | 0.05 - 0.15 | <0.001 | Women | -0.01 | -0.11 - 0.08 | 0.80 |
| Adjusted for gender, age, age<sup>2</sup>, first ten principal components. Associations are per SD change in PGS |  |  |  |  |  |  |  |  |

### Robustness checks

Robustness checks using ivreg2 confirmed the PGS was not a weak instrument. In UKHLS, the Cragg-Donald Wald F statistics (for all participants, men and women respectively) for analyses of own earnings were 303.7, 149.3, and 162.5, and for analyses of partners earnings 227.9, 125.3 and 112.2. In the smaller ELSA sample, they were for own earnings 79.5, 33.0, 47.0 and for partners earnings 66.0, 39.1, 28.8. Associations in UKHLS with gross earnings were similar to associations with net earnings (Supplementary Table S1). Methods based on individual SNP-level associations (IVW, MR-Median and MR-Egger regression) require more power than when using a polygenic score, but results were largely consistent with main results. MR-Egger regression found no evidence of directional pleiotropy (Supplementary Table 4), except for with odds of holding a professional or managerial occupation in UKHLS for men (Egger constant=-0.02, p=0.04) and all participants (Egger constant=−0.02, p=0.02) and in ELSA for women (Egger constant= −0.05, p=0.02).

## Discussion

This analysis used national samples representative of their target populations, and specifically considers labour income. Results are consistent with a causal association in men as well as women of BMI on own earnings, and of probability of employment, of having a university degree and of holding a professional or managerial occupation. Robustness checks indicated that for occupational social class, results may have been influenced by a degree of pleiotropy. There was suggestive evidence of an impact on earnings of a partner, although results were imprecise. Results do not support an impact on probability of partnership. These results are broadly consistent with the later UK Biobank paper(11), although the limited income data in that survey precludes direct comparison of models involving earnings. This supports generalizability of those results, despite the limited socioeconomic representativeness of the UK Biobank sample (for example, around 47%(11) of participants were university-educated, while the 2011 census found that 27% of people in England and Wales aged >16y had a degree, lower in the equivalent age range(27)). We add to previous work by exploring independent associations of genetically-instrumented BMI with own and partners’ earnings. Results suggest an additional link between own BMI and partners’ earnings, but imprecise estimates mean firm conclusions cannot be drawn.

In terms of mechanisms, our results find evidence for an impact of BMI on not just earnings but labour market participation (logistic models for whether in work), at odds with findings from UK Biobank(11). However, analysis of employment status was there restricted to participants in the labour market - and therefore comparing employed participants to jobseekers specifically, with economically inactive participants excluded. The discrepancy may therefore suggest any BMI-based selection has a greater impact on exit from the labour market altogether than from employment to unemployment. This is consistent with observational evidence for differential health-based selection into unemployed and economically inactive groups(28).

We found the weighted polygenic score for BMI significantly predicted self-rated health for UKHLS and ELSA participants. This suggests part of the causal impact of BMI on labour market outcomes may operate via health, consistent with the substantial attenuation of OLS estimates seen when self-rated health was adjusted for. However, since impaired self-rated health may be both a cause and a result of labour market adversity(29), the mechanisms linking genetically-determined BMI, labour market success and self-rated health are likely to be complex. Meanwhile, the evidence for an impact of genetically-instrumented BMI on educational qualifications, here captured by likelihood of holding an undergraduate degree, suggests effects could be partially mediated by educational attainment. Other mechanisms such as weight-based discrimination in the workplace likely play a role but could not be investigated here.

Larger estimates for the impact of BMI on own earnings in the IV analysis than adjusted OLS models were unexpected, and suggest several possible explanations. One is that associations in observational models may be suppressed by confounding, which could happen if low income can, under certain circumstances, lead to lower body weight. This would be consistent with a previously documented association between long-term unemployment and reduced odds of overweight in UKHLS(30). A second explanation relates to the fact that earnings are plausibly a function of mean BMI over a lifetime, which may be better indexed by the externally-weighted PGS than by a single observed BMI measurement. This is supported by evidence showing the influence of gene variants on BMI emerges in adolescence and remains throughout adult life(31). Beta coefficients for BMI-SNP associations from GWAS are themselves estimated using single BMI measurements in large but finite samples, and therefore subject to noise, but nevertheless relate specifically to a very stable component of inter-individual variation in BMI. BMI-earnings associations from IV models may therefore be less influenced by downward bias due to classical measurement error. Thirdly, IV results may have been inflated by a degree of pleiotropy, which the robustness checks performed may have been underpowered to detect. Finally, estimates from Mendelian Randomization relating to social outcomes may be biased by two further processes whose investigation requires genetic data on related individuals(32), not available in these surveys. Dynastic effects - an influence of parental genotype on offspring phenotype via the offspring’s environment, sometimes called ‘genetic nurture’ - can inflate estimates based on inherited genotypes(33). A second issue is assortative mating - that people tend choose partners who are more similar to themselves than would be expected by chance. This can bias associations between genetic variants for one trait and an outcome phenotype(34). Since results from our analyses of partners supports a degree of cross-trait assortative mating between BMI and earnings, and with evidence of an associations between partners’ polygenic scores in one survey, it is certainly conceivable that assortative mating may have influenced results regarding own labour market outcomes. Further work in large surveys with genetic information across families will be required to fully separate these mechanisms.

## Strengths and limitations

A major strength of this analysis is the socioeconomic representativeness of the samples, and replication in a second study population where exposure was assessed using the same method and data collection agency. The high-quality income data meant phenotype was very well assessed in comparison to previous studies, and because income was collected for individuals rather than just households, we were able to examine relationships within couples. A limitation is the comparatively small individual sample sizes for a genetic analysis, which led to imprecise estimates of study-specific relationships especially in ELSA. The relationship of BMI with both health(35) and aspects of socioeconomic position(30, 36) may not be linear, suggesting causal influence of BMI on earnings may differ along the BMI spectrum. While methods for non-linear IV analysis exist and have been applied to similar questions(11), we were unable to apply them due to sample size limitations(37). A degree of assortative mating, indicated by results of partners’ analyses, could have influenced other results. Finally, as with any MR study, an influence of pleiotropy cannot be ruled out.

## Conclusion

In two nationally-representative samples of UK adults, genetically-instrumented BMI was associated with lower earnings, probability of being in work, of holding a managerial or professional occupation, and of having a university degree, but not with partnership. Results broadly support previous work suggesting an influence of BMI on diverse aspects of socioeconomic position, indicating that targeting inequalities in BMI may also act on inequalities more widely.

## Legend for Figures 1–6

*First three clusters:* OLS estimates adjusted for age, age^2^, gender, the first ten principal components

*Second three clusters:* OLS estimates adjusted for age, age^2^, gender, Government Office Region, smoking status, self-rated health, educational qualifications (except for odds of holding a university degree) and the first ten principal components.

*Third three clusters:* IV estimates adjusted for age, age^2^, gender, the first ten principal components

